# DeepProjection: Rapid and structure-specific projections of tissue sheets embedded in 3D microscopy stacks using deep learning

**DOI:** 10.1101/2021.11.17.468809

**Authors:** Daniel Haertter, Xiaolei Wang, Stephanie M. Fogerson, Nitya Ramkumar, Janice M. Crawford, Kenneth D. Poss, Stefano Di Talia, Daniel P. Kiehart, Christoph F. Schmidt

## Abstract

The efficient extraction of local high-resolution content from massive amounts of imaging data remains a serious and unsolved problem in studies of complex biological tissues. Here we present DeepProjection, a trainable projection algorithm based on deep learning. This algorithm rapidly and robustly extracts image content contained in curved manifolds from time-lapse recorded 3D image stacks by binary masking of background content, stack by stack. The masks calculated for a given movie, when predicted, e.g., on fluorescent cell boundaries on one channel, can subsequently be applied to project other fluorescent channels from the same manifold. We apply DeepProjection to follow the dynamic movements of 2D-tissue sheets in embryonic development. We show that we can selectively project the amnioserosa cell sheet during dorsal closure in *Drosophila melanogaster* embryos and the periderm layer in the elongating zebrafish embryo while masking highly fluorescent out-of-plane artifacts.

## Introduction

Three-dimensional fluorescence microscopy of transparent model organisms such as the fruit fly *Drosophila melanogaster*, the nematode *Caenorhabditis elegans* or the zebrafish *Danio rerio* is a central tool in developmental biology. With modern techniques, the dynamics of entire organisms can be rapidly imaged as sequences of stacks of two-dimensional image slices, resulting in GBs of data per recording. Manual image processing is far too slow to mine such data, and potentially introduces bias. Convolutional neural networks, a form of machine learning, have been shown to far outperform conventional algorithms for visual feature extraction in many areas of research and engineering, including the life-sciences ^1^. Neural networks such as U-Net ^2^ can robustly segment and classify complex image features, defined by an initial training process.

We here present a new deep learning approach that allows us to automatically extract image content from dynamic curved 2D manifolds embedded in 3D image stacks of developing tissues. DeepProjection (DP) is an algorithm for structure-specific, surface projections based on feature detection with convolutional neural networks. DP is a useful tool for developmental biology because, throughout phylogeny, cell sheet migration is a fundamental feature of morphogenesis in development and in wound healing. Cell sheets assume complex curved geometries and move to form internal organs and structures, including the neural tube, the guts and the heart in vertebrates. To follow tissue sheet morphogenesis in living embryos, it is necessary to extract selected sections of image content from many slices of each 3D stack to create 2D projections, while rejecting content from other planes. Past approaches have severe shortcomings. Maximum intensity projection (MIP) is simple and fast, but only works when fluorescence from the structures of interest dominates noise and off-target labels. MIP cannot differentiate subtle and low-intensity content of interest from bright content in other image planes. Manual omission of problematic slices or reduction of the overall imaging volume can also eliminate parts of the target content. Manual masking of individual slices is entirely impractical for long recordings. Other reported methods for z-projection use pixel value statistics, *e*.*g*., the minimum, the median or the sum of the pixels along the z axis through the stacks, typically assuming that the structures of interest display the brightest fluorescence or have the sharpest contrast, but don’t use information from neighboring pixels. The extracted 2D manifolds are easily distorted by single bright pixels, *i*.*e*., the masks are not continuous, have holes and no sharp edges. The extracted 2D manifolds are therefore not smooth. More sophisticated approaches can be classified into three categories: (i) smoothing of height maps derived by MIP ^3^, (ii) ranking of z-slices by visual pattern recognition of targeted tissue structures by edge filters ^4^, Fourier transforms or wavelet transforms ^5^, or (iii) evaluating mean and variance of intensity distributions in the neighborhood of each pixel ^6^. All these algorithms only perform well when the target structures are bright and clearly distinguishable from background noise. Bright fluorescent structures with a clear texture in the image background, such as auto-fluorescent yolk granules in *Drosophila* embryos, are not robustly discriminated against ^7^. Most importantly, all these approaches are static, *i*.*e*. require manual parameter optimization for each new recording. Machine learning approaches, in contrast, can be trained to deal with a broad range of imaging conditions. One pioneering application of ML to developmental imaging is CSBDeep, a package for content-aware image restoration ^8^. This package demonstrates how convolutional neural networks can be used for combined projection and denoising of microscopy stacks of *Drosophila* wing development, by coupling a small convolutional network for 2D projection and a U-Net for subsequent denoising ^2, 8^. However, due to the strong emphasis on denoising, original pixel intensities are not conserved in the output, and the algorithm does not yield the manifold containing the tissue.

The key unaddressed challenge is to design an automated approach that can (i) detect defined fluorescent features that do not solely stand out by intensity or texture, and (ii) project the entire 2D manifold that the detected features reside in without distorting intensities while completely rejecting content from regions outside of this 2D-manifold. The critical advantage of convolutional neural networks is that they can be trained to simultaneously detect various distinctive features of the target structure and then also select image content based on, but not limited to the detected structures.

DP uses an encoder-decoder neural network that analyzes features and textures in a 3D stack to create binary masks which contain only the areas specified as targets in the training data, in our case tissue layers containing crisp cell boundaries. Thereby DP omits out-of-plane fluorescent structures and artifacts. We demonstrate DP using two models, dorsal closure in fruit fly (*Drosophila melanogaster)* development ^9, 10^ and periderm development in zebrafish (*Danio rerio)* embryogenesis ^11, 12^. DP predicts a single stack in just 1-10 s, depending on stack size. Significant gains in signal-to-noise ratio allow us to resolve even subtle structures. DP yields time consistent results for time-lapsed movies since the algorithm detects persistent spatially extended features in the target manifold and is not deflected by temporary fluctuations and artefacts. DP produces masks that select 2D manifolds without modification of intensities in those planes. We show how these masks can be used to select image content from other fluorescent channels from the same 2D manifolds by extracting actin dynamics near the apical surface of amnioserosa cells during dorsal closure of *Drosophila*. We further show how the detected 3D geometry can be used to unfold curved cell sheets.

## Results

We developed DP to be broadly applicable for the rapid, automated processing of time-lapsed 3D recordings of developing embryos and tested it on recordings of dorsal closure in *Drosophila* embryos and periderm development in zebrafish embryos. DP is a custom-designed convolutional neural network that locally analyzes the image stack and classifies complex morphologies and textures. DP was trained with pairs of image stacks and the corresponding manually created binary masks. DP can simultaneously detect various target structures defined by the user through training, and decide on which regions of a given slice to keep. In our applications to epithelial tissue sheets, *e*.*g*., only the regions of the stack containing tissue with crisp cell boundaries are retained (Figures 1A-H). Dorsal closure during *Drosophila* embryogenesis (Fig. 1A, C, E, F) occurs 12-15 hrs after egg laying ^10, 13^. At this stage of development, the dorsal opening, left behind after germ band retraction, is covered by a curved sheet of squameous epithelial cells (amnioserosa (AS) cells). The dorsal opening closes in ∼3hrs while AS cells subduct under the lateral epidermis (LE) or apoptose ^13^. We labeled cell boundaries by E-cadherin-GFP. Analyzing AS dynamics was complicated by the curved shape of the AS and highly fluorescent yolk granules and subducted cells underneath the AS tissue (Fig. 1A, E). Tracking LE cells was impeded by low signal-to-noise ratios. DP clearly resolved AS and LE cell boundaries while yolk particles and gut cells were masked (Fig. 1C, E, F). During zebrafish development (Fig. 1 B, D, G, H), the periderm covers the entire elongating embryo (Fig. 1B’). As the embryo narrows towards the posterior region, the left and right sides appear overlaid in the MIPs (Fig. 1B) when imaged laterally. This makes it impossible to distinguish the upper from the lower tissue layer. Further, due to the strong curvature, projected cell shapes and areas are distorted. DP distinguished the tissue of interest from lower layers and the notochord, revealing the cell boundaries and making it possible to study cell shapes (Figures 1D, G, H).

**Figure 1:**
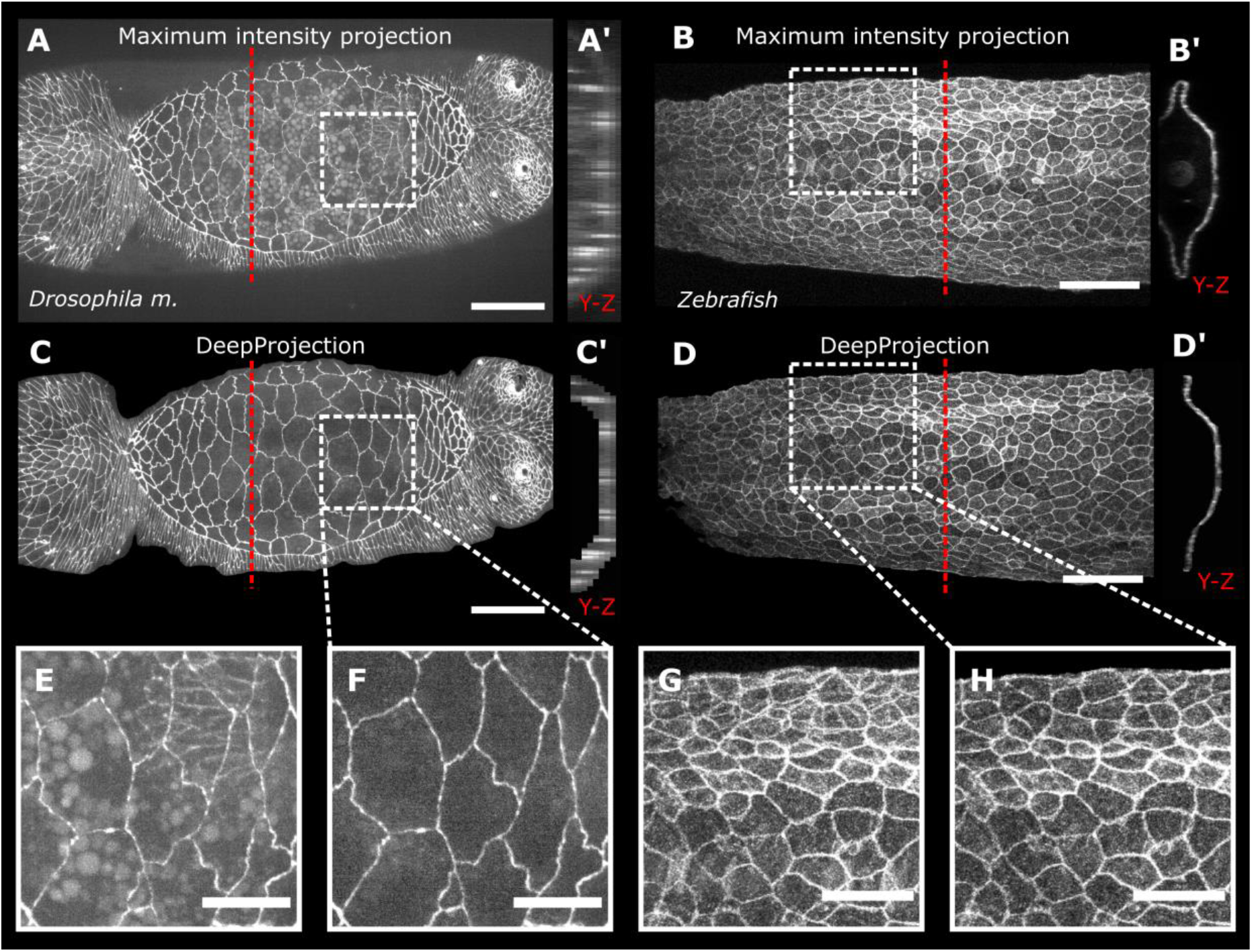
Comparison of DeepProjection (DP) with maximum intensity projection (MIP). **A:** MIP of a single stack (8 slices, 1 μm z-distance) of images of the dorsal opening of a *Drosophila* embryo during dorsal closure, cell boundaries labeled with Cadherin-GFP. **B:** MIP of as single stack (53 slices, 2 μm z-distance) of images of zebrafish periderm labeled by *krt4*-directed lyn-EGFP fluorescence 1 day post fertilization (dpf). **A’-D’:** y-z cuts of 3D image stacks at red dashed line in A-B. **C’** and **D’** show the masked stack with the manifolds predicted by DP. **C, D:** DP results from the same stacks of *Drosophila* and zebrafish embryo. **E, F:** Zoom into amnioserosal tissue in *Drosophila* comparing MIP (**E**) and DP (**F**), showing successful masking of yolk granules and gut tissue underneath the amnioserosal tissue. **G, H:** Zoom into zebrafish embryo comparing MIP (**G**) and DP (**H**), showing masking of underlying epithelial tissue layer. **Scale bars:** A, C 50 μm; B, D 100 μm; E-F 10 μm; G, H: 50 μm.

DP analyses 3D microscopy stacks (Fig. 2A) using an encoder-decoder convolutional neural network (Fig. 2B), inspired by the U-Net architecture ^2^. The left branch of the neural network extracts high-dimensional features by pairs of 3D convolutions with kernel size 3×3×3, each followed by Scaled Exponential Linear Units (SELU) activation. Between each double-convolutional layer, the content is down-sampled by max-pooling with kernel size 1×2×2 in only the x and y directions. The number of convolution kernels further doubles in each layer. This multi-layer structure ensures the efficient extraction of both high-frequency features (such as bright cell boundary pixels) and low-frequency image features (such as whole cells in a tissue context). After feature extraction, the feature map is decoded and up-scaled again to the initial stack dimensions using up-convolutions with kernel size 1×2×2 alternating with 3D convolutions with kernel size 3×3×3. To preserve the spatial resolution of shapes and boundaries, the input of each decoding layer is concatenated with the output of the corresponding encoding layer. After the last up-convolution layer, the output is scaled between 0 and 1 with sigmoid activation, yielding binary masks (Fig. 2C). In a last step, the predicted masks are multiplied with the input stack, creating a masked stack with the target structure, in the case shown tissue with crisp cell boundaries (Fig. 2D). A maximum z-projection inside the predicted manifold then yields the final DP result (Fig. 2E). The predicted masks for each stack can be exported and subsequently applied to other fluorescent channels.

**Figure 2:**
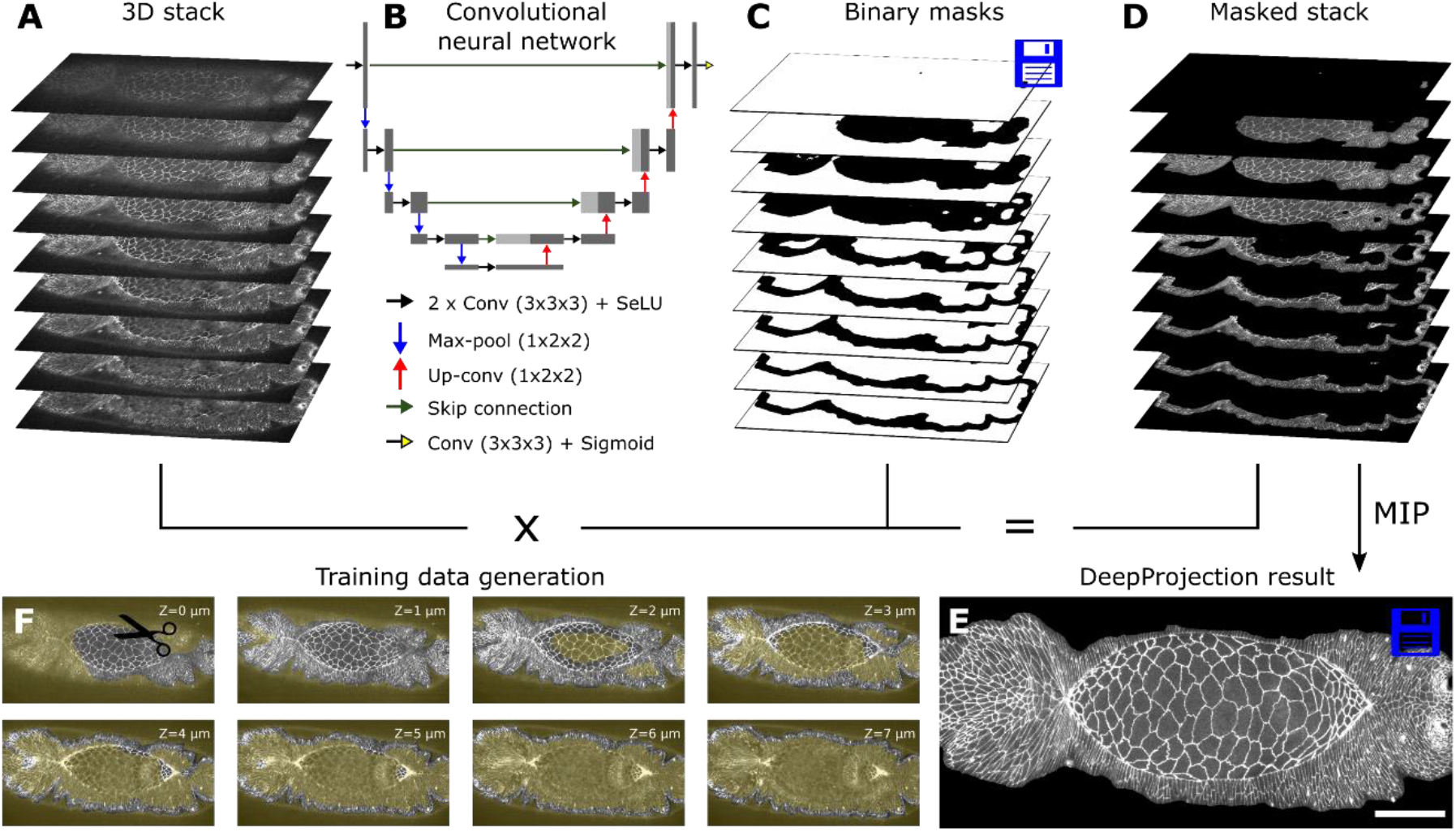
DeepProjection algorithm. **A:** Input 3D stack of *Drosophila* dorsal closure. **B:** Convolutional neural network architecture. **C:** Output of neural network showing binary masks and mask edges. **D:** Multiplication of input stack with predicted binary masks yields masked 3D stack containing only target tissue with crisp cell boundaries. The predicted masks can be saved and applied to other fluorescent channels. **E**: MIP of masked 3D stack yields the result. **F:** Illustration of manual training data generation. Yellow areas are cut out manually for each slide and only areas containing the target tissue are kept. **Scale bar:** E 50 μm.

The ground truth (GT) used for training was created manually by human experts cutting out unwanted content (yolk granules, noise background, blurry out-of-focus cell boundaries, off-target tissue layers) of each image slice and keeping only the structure of interest (target epithelial tissue with crisp tissue boundaries) with the freehand selection tool in *Fiji/ImageJ* (Schindelin et al., 2012) (Fig. 2D). We selected not just the cell boundaries, but whole cells. The masked stacks were binarized by clipping all remaining parts to 1. The generation of 100 masked stacks for training took ∼15 hours. This step only needs to be performed once. Our training data contained stacks of varying image quality and three different labeling strategies, with different image resolution, recorded with different microscopes (100 stacks for *Drosophila*, 20 stacks for zebrafish). We augmented the training data 6-fold by randomly flipping and adjusting brightness and contrast. We aimed for fully binary masks with sharp and straight edges and contiguous areas. For training, we chose a log-cosh-Tversky loss function that yields sharp and crisp mask edges ^14^. This loss function can be tuned to punish false positives (high α, low β) or false negatives (low α, high β), and is identical to the common Dice loss for α = β = 0.5. We found optimal training results for α = 0.3 and β = 0.7. We trained two separate networks, one for *Drosophila* and one for zebrafish, in each case for 50 epochs with learning rate 1e-5, stack patches of (512×512) pixels in x, y and batch size 12 on a workstation with an NVidia GTX 1080 Ti GPU.

To demonstrate the capabilities of DP, we compared its performance with simple MIP and three published algorithms FastSME (FSME) ^15^, Local Z Projector (LZP) ^6^ and CSBDeep (CSBD) ^8^. We trained CSBDeep with the MIP of the masked stacks of the DP training data as GT (using default parameters, 200 epochs with learning rate 4e-5). We selected representative confocal stacks from recordings of *Drosophila* dorsal closure (N = 8) and zebrafish periderm (N = 4), distinct from the training data, and compared the respective results with manually-created GT (Fig. 3A-C). The parameters of FSME and LZP were optimized individually for each stack. For the dorsal closure stacks, DP was able to reproduce the ground truth, while MIP and FSME failed to remove yolk granules and underlying gut tissue; LZP did detect only parts of the AS tissue and yolk granules leaked through; CSBD showed holes and yolk granules in the cell centers and the intensity grey values appeared distorted (Fig. 3A). The faint cell boundaries of the lateral epidermis tissue were well detected by DP and LZP, while FSME did not perform better than MIP. CSBD showed opaque, fog-like artifacts (Fig. 3B). Subducting cells along the seam under the LE were only discriminated against by CSBD and DP. CSDB distorted intensity values (Fig. 3B). For the zebrafish periderm, DP, LZP and CSBD were able to differentiate upper from lower tissue layers, unlike MIP and FSME (Fig. 3C). However, FSME, LZP and CSBD produced artifacts as black lines and cell boundary snippets, due to the high local gradient of the tissue at the edge (Fig. 3C). CSBD again distorted intensity values, and high-frequency details inside the tissue manifold appeared smoothed or were missing (Fig. 3C). The CSBD algorithm attenuates or emphasizes certain features which is evident in a plot of pixel grey values of the results against the maximal intensity projection (Fig. 3D). FSME, LZP and DP yield pixel grey values below the MIP result, as expected for unique z-maps and binary masks. CSBD, however, nonlinearly distorts the pixel values, which makes it impossible to quantitate protein concentrations (Fig. 3D).

**Figure 3:**
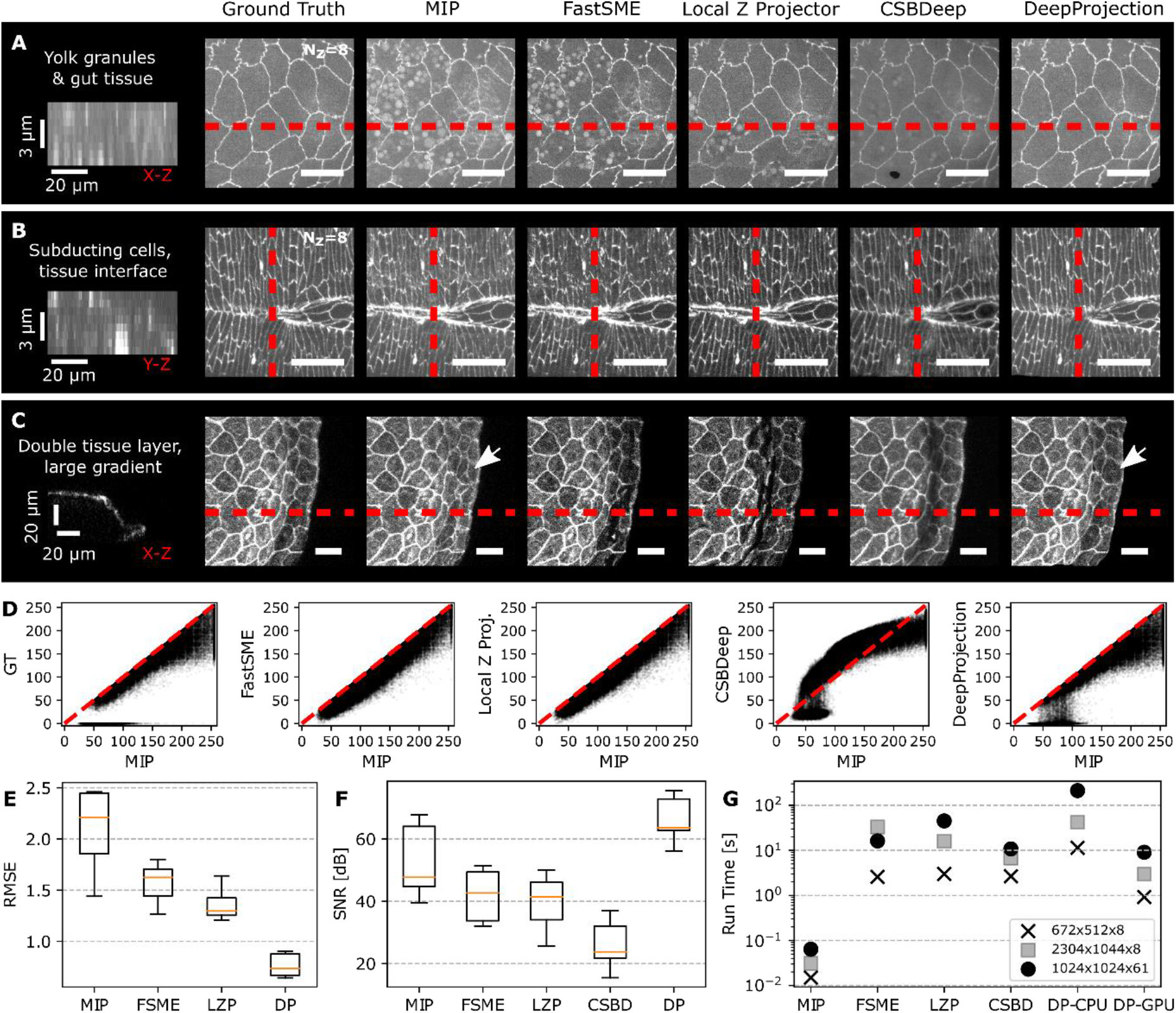
Comparison of DP with published algorithms. **A:** Results for a single confocal stack of amnioserosa (AS) tissue during early *Drosophila* dorsal closure, with auto-fluorescent yolk granules and gut cells. **B:** Results for amnioserosa-lateral epidermis (LE) tissue interface and canthi during late *Drosophila* dorsal closure, with faint tissue cell boundaries, subducting cells and interface of two different tissue types. **C:** Results for zebrafish periderm (1dpf), with large tissue gradient and second tissue layer underneath. Images on the left of **A-C** show vertical cuts at red dashed lines, each averaged over 10 pixels perpendicular to line. **D:** Pixel-wise scatter plot of algorithms against MIP to check for distortion of fluorescent grey value. **E:** Root-Mean-Square errors of algorithm results relative to ground truth. **F:** Signal-to-noise ratio of algorithm results with ground truth as reference. **G:** Log-scale plot of algorithm run time for three different stack sizes. In box-and-whisker plots, boxes show median (red) with IQR, with whiskers extending to the 5^th^ and 95^th^ percentile. **Scale bars:** 20 μm.

We further evaluated the performance of all methods in three ways: (i) We calculated the normalized root-mean-square errors (RMSE) of the z-maps *z*(*x,y*) with respect to the ground truth, the z-index of the selected pixel:

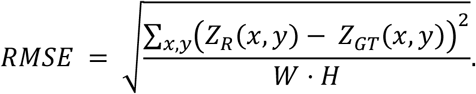

Since DP yields masks with potentially more than one slice selected per x-y pixel, we created unique z-maps for DP and ground truth by selecting the z-index corresponding to the maximum intensity inside the manifold. The CSBD package does not output a z-map or binary masks. DP strongly outperformed all other algorithms (Fig. 3D). (ii) We calculated the signal-to-noise ratio (SNR) of the reconstruction results *I*_*R*_ with respect to the ground truth *I*_*GT*_:

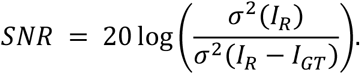

DP performed significantly better than both FSME and LZP (Fig. 3E). Interestingly, even MIP performed better than FSME and LZP, since all bright features, desired and undesired ones, are conserved by MIP. FSME and LZP both create a smooth z-map with only one selected plane per x-y pixel, whereas DP predicts a set of binary masks embedding the tissue. When more than one slice is selected for a given x, y pixel, DP then uses MIP inside this embedding to get the final 2D projection result. When the structures of interest (in this case fluorescent cell boundaries) span multiple z-slices, MIP and foremost DP yield better SNR results by including relevant signal from more than one slice. Due to the non-linear distortion of grey values, CSBD scored poorly (Fig. 3D). (iii) We assessed the run time of algorithms on exemplary stacks with three different sizes (Fig. 3F). MIP was very fast due its simplicity. DP required about 1 s to predict a stack with dimension 8×640×512 pixels, when run on a graphics card (GPU), 3x faster than FSME and LZP. When run on the CPU, DP prediction took around 10x longer than on the graphics card, but was still within a practical range. CSDB run times on a GPU were slightly longer than DP.

To demonstrate the option of mask transfer to other simultaneously recorded fluorescent channels, we performed dual-color imaging of dorsal closure with E-cadherin-tomato and GFP-actin. The MIP of the actin channel shows intracellular actin networks and actin-rich filopodial cores (Fig. 4A). However, it is not possible to distinguish between apical actin, responsible for the contraction of apical cell areas ^16, 17^, and actin elsewhere in the cells. We next used DP to predict binary masks using the cell boundary information captured in the E-cadherin channel and then applied the masks to the actin channel (Fig. 4B, D). As highlighted in figure 4B’, the predicted masks include the apical surface with actin networks and filopodia. A comparison between the DP and MIP results shows additional pronounced actin structures at the basal surface of cells visible in MIP, but not in DP (Fig. 4A’,B’). In order to extract the basal actin, we shifted the binary masks by 4 pixels (corresponding to 2 μm in the z-direction) (Fig. 4E), and then applied them to the actin channel (Fig. 4C). DP thus allows us to not only extract image content from other channels from the predicted 2D manifolds, but also from other planes, offset in the z-direction.

**Figure 4:**
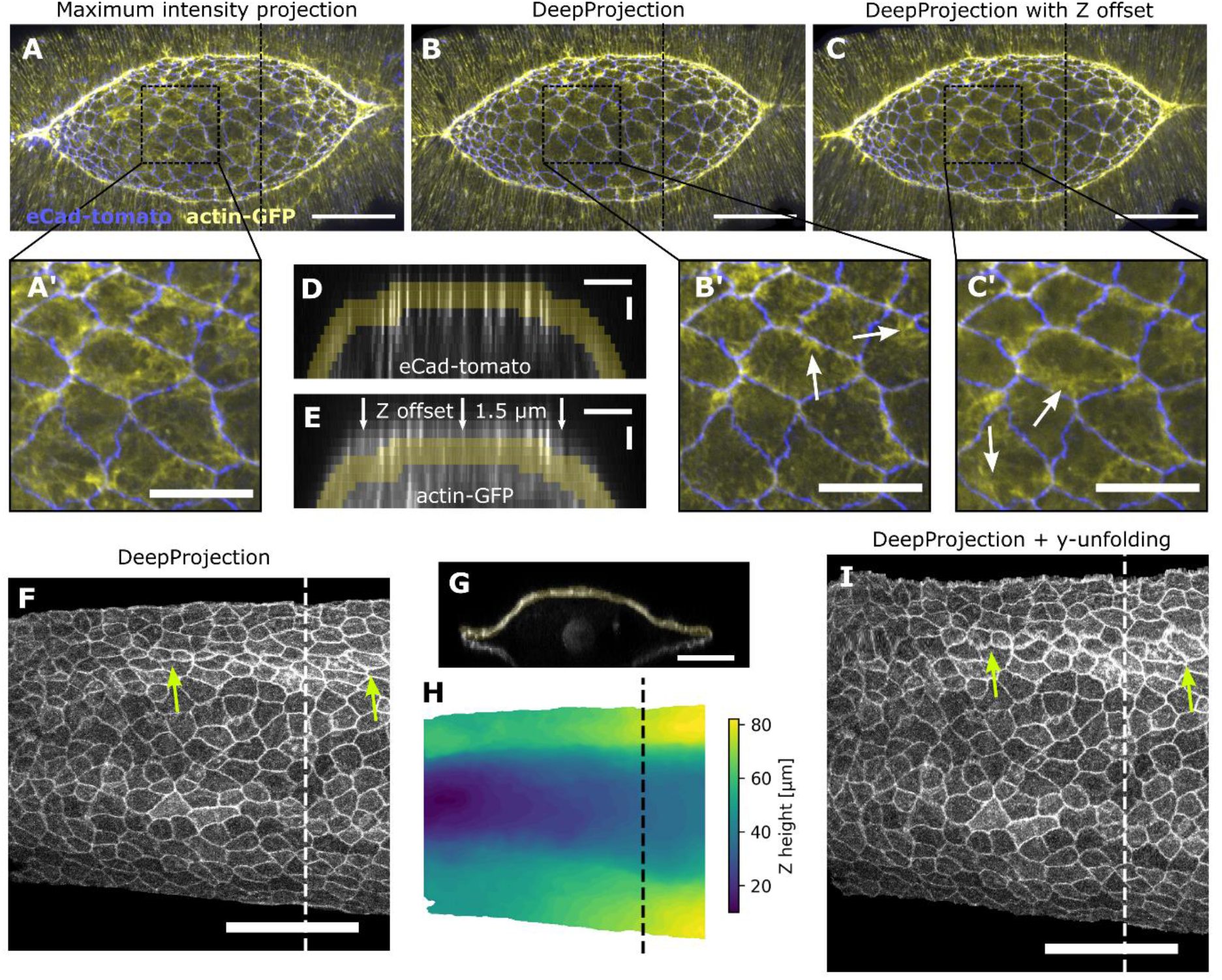
Mask transfer to project content from other fluorescent channels. **A:** MIP of dual-color confocal stack of *Drosophila* embryo during mid-stage of dorsal closure with labeling via E-cadherin-tomato and actin-GFP. **B:** Result of applying the masks, predicted by DP from E-cadherin cell boundaries to the actin channel, showing mainly actin-rich filopodia at the apical surface of cells. **C:** Applying the same masks with a Z-offset of 4 pixels/2 μm to the actin channel, showing contractile actin networks underneath the apical surface. Scale bars: A-C 50 μm, A’-C’ 20 μm. **D:** Y-Z cut of E-cadherin-tomato channel (at dashed line in A-C) with binary mask predicted by DP. **E:** Y-Z cut of actin channel with binary masks with Z-offset. **F:** DP result of zebrafish periderm development (1dpf). **G:** x-z cut along white dashed line in **F**. The manifold predicted by DP is highlighted in yellow. **H:** z-map calculated by averaging the manifolds’ positive indices at each x-y position. The z-map was subsequently smoothed with a mean filter with kernel size 5×5 pixels. **I:** Result of unfolding algorithm applied in y direction. As highlighted, the unfolding reveals the true shape and area of cells at positions with large gradient. **Scale bars:** D, E 20 μm in y, 1.5 μm in z; F, I 50 μm.

If a curved cell sheet displays steep gradients, cell shapes and areas are distorted in z-projections (Fig. 4F, G). To unfold such tissue sheets, we created unique z-maps by averaging the z-indices with value 1 of the binary masks predicted by DP, and then smoothed the z-maps by 2D averaging with kernel size of 3×3 pixels (Fig. 4H). The gradient tensor *α*(*i,j*) of the z-maps then yields the local curvature in x and y. This makes it possible to locally correct the distortion for individual cells using an affine transformation with a gradient tensor of the z-map averaged over the close proximity of each cell (not shown here). Alternatively, we can unfold the tissue in just one direction. This is particularly useful for tube-like tissues. First we define a marker line at position *k*. Then we cut the curved tissue in 1 pixel wide stripes, straighten them, stitch them back together and align the stripes at the previously defined marker line *k*. The unidirectional transformation map *T*(*i,j*) for each pixel (*i,j*) can thus be defined as:

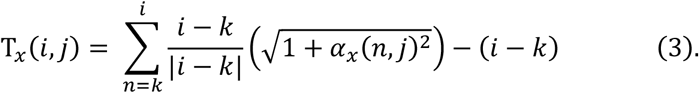

The unfolding algorithm successfully straightens the regions of the zebrafish embryo with high gradients in x, revealing the true cell sizes and shapes (Fig. 4I). Our approach is similar to previously reported unrolling and cartographic approaches ^18^.

So far we have focused on the projection of individual confocal stacks. When evaluating time-lapse recordings of developing embryos, time consistency of the projection method becomes important. We applied DP to time-lapse recordings of dorsal closure (Supplementary movie M1). Even though the stacks were predicted one-by-one, and no information was propagated between consecutive time points, the results were time consistent, demonstrated by the stable position of tissue edges (Movie S1). DP robustly detects time persistent image features, of both low and high spatial frequencies and is thus not deflected by time variant noise or decreasing fluorescent intensity due to bleaching.

## Conclusions

DP uses a convolutional neural network to selectively extract and project image content from curved 2D manifolds embedded in 3D confocal stacks. DP detects complex features that are in each application specified by user-annotated training, which makes it possible to mask even highly fluorescent artifacts while faithfully detecting weakly fluorescent structures of interest. DP could extract and project image content from dynamic curved tissue sheets in both *Drosophila* and zebrafish embryos while masking background content and noise. Image processing with DP greatly simplifies the segmentation and tracking of individual cells in subsequent processing steps. Original fluorescence intensity values in the selected manifold are strictly preserved for quantitative analysis. DP significantly outperformed the alternative algorithms we tested and created time-consistent results for time-lapsed recordings. DP is substantially faster than most published algorithms. Due to its universal architecture, DP is not limited to the analysis of epithelial tissues, but can be applied to extract any 2D manifold from 3D data when properly trained for the features of the target manifold. For high-content imaging pipelines, DP can rapidly and robustly compress data from stacks to single images to save storage space without losing information from the target manifolds. Deep learning algorithms such as DeepProjection constitute a major leap in the capability to process massive imaging data and will enable researchers to rapidly mine data and rigorously quantify complex morphogenetic processes.

## Methods

### Preparation and imaging of *Drosophila* embryos

Cell junctions were labeled with either DE-cadherin-GFP or DE-cadherin-mTomato (labeling *Drosophila* E-cadherin), both knock-in lines under control of the endogenous promoter ^19^. F-actin was labeled with the GFP-moesin actin binding domain, expressed under the control of the *spaghetti squash* promoter in the sGMCA line ^9^. All stocks were maintained at room temperature or 25°C on standard cornmeal/molasses fly food or in embryo collection cages with a grape juice agar plate and yeast paste. Embryos were collected either 2–4 hours after egg lays and aged overnight at 16°C, or from overnight egg lays at 25°C. To remove the chorion, embryos were incubated in a 50% bleach solution for 1.25 min and then rinsed extensively with deionized water. Pre-dorsal closure stage embryos were selected using a reflected-light dissecting microscope. Embryos were prepared for imaging as described previously ^20^. Images were acquired using Micro-Manager 2.0 software (Open Imaging, San Francisco, CA) to control a Zeiss Axiovert 200M microscope (Carl Zeiss, Thornwood, NY) equipped with a Yokogawa CSU-W1 spinning disk confocal head (Solamere Technology Group, Salt Lake City, UT), a Hamamatsu Orca Fusion BT camera (Hamamatsu, Japan), and a Zeiss 40X LD LCI Plan-Apochromat 1.2NA multi-immersion objective (glycerin). Due to the curvature of the embryo, we imaged multiple z planes for each embryo at each time point to view the dorsal opening. We recorded 8 z-slice stacks with 1 μm step size for single color, and 14 z-slice stacks with 0.5 μm step size for dual-color movies. Stacks were acquired every 15 s throughout the duration of closure with a 100 ms exposure per slice for GFP and a 150 ms exposure per slice for mTomato.

### Zebrafish husbandry and sample preparation for live imaging

Zebrafish of the Ekkwill strain were maintained between 26 and 28.5 °C with a 14:10 hour light:dark cycle. Fish between 3-6 months were used for experiments. Transgenic *krt4:lyn-EGFP* fish were described previously ^21^. Male and female fish were set up for mating in tanks with dividers. The dividers were removed in the morning for timed mating. Embryos were collected in E3 medium and screened at 1 dpf for expression of GFP in the periderm (krt4-lynGFP). The positive embryos were transferred to a dish with E3 medium and tricaine (Sigma E10521-50G) at 0.01% concentration. The embryos were dechorionated with forceps and mounted in fluorinated ethylene propylene (FEP) tubes according to published protocols ^22^. The FEP tubes were coated with 3% methlycellulose and embryos were mounted in 0.1% agarose with 0.01% tricaine to immobilize them during imaging. The tube was then placed in a 60 mm culture dish with an agarose bed, held in place with 1% agarose and immersed in E3 medium. Images were acquired with LASX software on an Leica SP8 confocal microscope using an HC Fluotar L 25X/0.95NA W VISIR water-immersion objective at 0.75 or 1X zoom. Image stacks with 40-60 slices were acquired every 15 minutes with a z-step size of 2 μm. Work with zebrafish was approved by the Institutional Animal Care and Use Committee at Duke University.

### DeepProjection software and hardware

DeepProjection is implemented in python 3.8 using standard free packages: numpy 1.19 for scientific computing, pytorch 1.7.1 (with cuda 11.0) for deep learning, albumentations 0.5.2 for data augmentation. The DeepProjection code was tested on Windows 10 and Ubuntu (Linux) 20.04 operating systems. The DeepProjection repository (https://github.com/danihae/DeepProjection/) contains the DP code and Jupyter notebooks for training and prediction with detailed instructions. There, we also provide pretrained network weights, training data and test data for *Drosophila* dorsal closure and zebrafish periderm. The DeepProjection package is further available on Python Package Index (PyPI). DeepProjection was trained on a workstation with 32 GB onboard memory and NVIDIA GeForce 1080 Ti. Prediction was successfully tested on regular laptop without dedicated GPU.

## Acknowledgements

We thank David Carlson, Duke University, for input during the algorithm design process. D.H. thanks the German Academic Foundation for funding during this work. D.P.K. acknowledges support from N.I.H. (R35GM127059), C.F.S. from the Soft Matter Center, Duke University. This was work was in part supported by an N.I.H. grant (R01-AR076342) to K.D.P and S.D.

## Author contributions

C.F.S., D.P.K., and D.H. designed the project; C.F.S. and D.P.K. supervised the project; D.H. and X.W. designed the algorithm, analyzed the results and compared DP with other algorithms; D.H. built the Python package and the graphical user interface; S.M.F. and J.M.C. produced the *Drosophila* data, N.R., K.D.P. and S.D.T. provided the zebrafish data; D.H., X.W., N.R. and S.M.F. created training data; D.H., C.F.S., and D.P.K. wrote the paper with input from all authors.

## Supplementary materials

**Figure S1:**
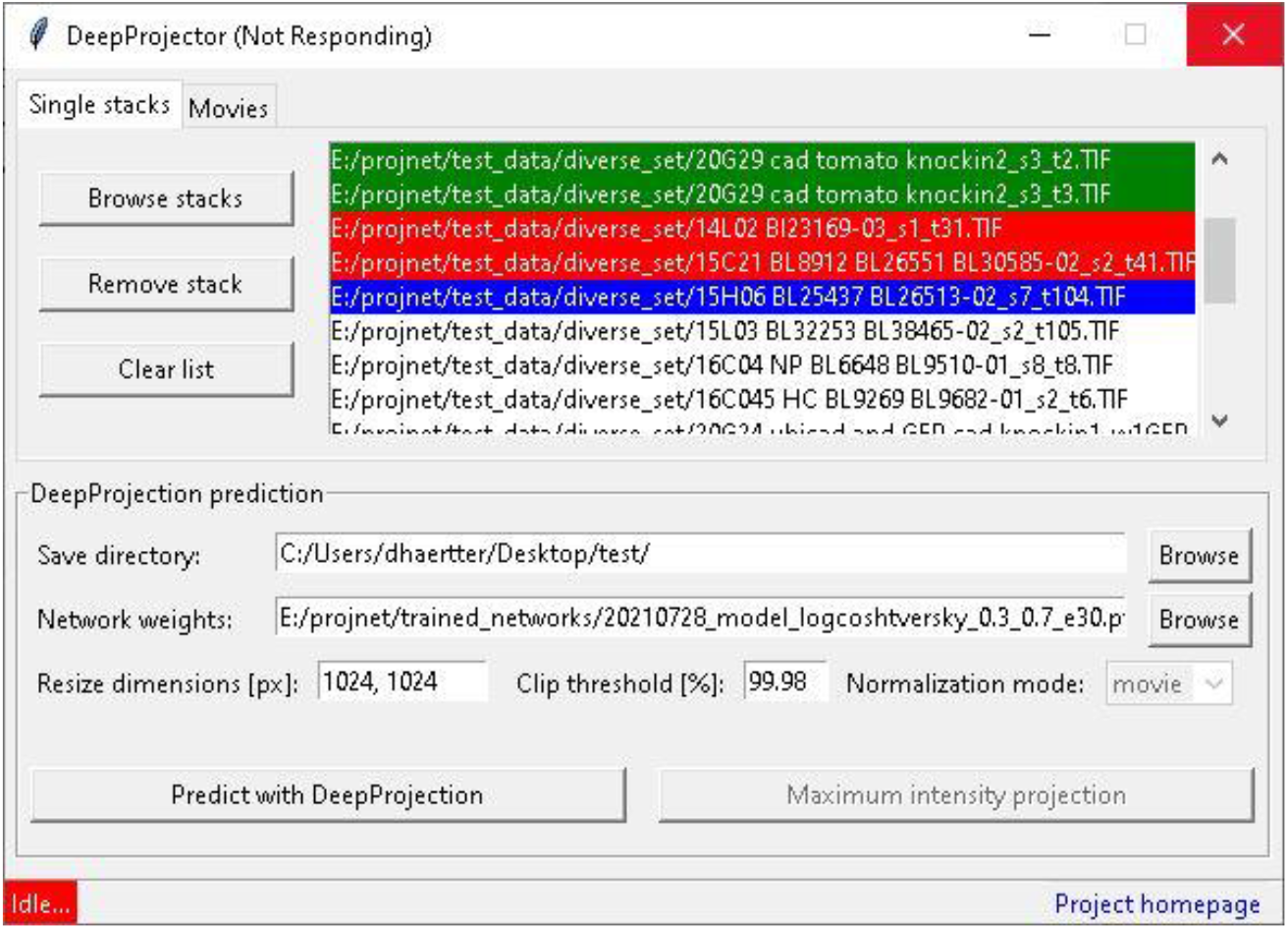
Graphical user interface (GUI) for prediction. At the top, multiple stacks or directories with time-lapse movies can be selected using the file browser. Single items can be added and deleted from the list. At the bottom, single stacks or movies in the list above can be predicted using DeepProjection. Parameters can be adjusted and the directory for saving the results and the trained neural network weights can be chosen. During prediction, progress is indicated by colored list items (green = successful, red = error, blue = in progress). Error events occur if input files are corrupted or the GPU memory or hard drive is full. Download: https://e.pcloud.link/publink/show?code=kZM7fJZOHPWjuH9iUpz8fS8uIp2OmHMwRyy

**Movie M1:**
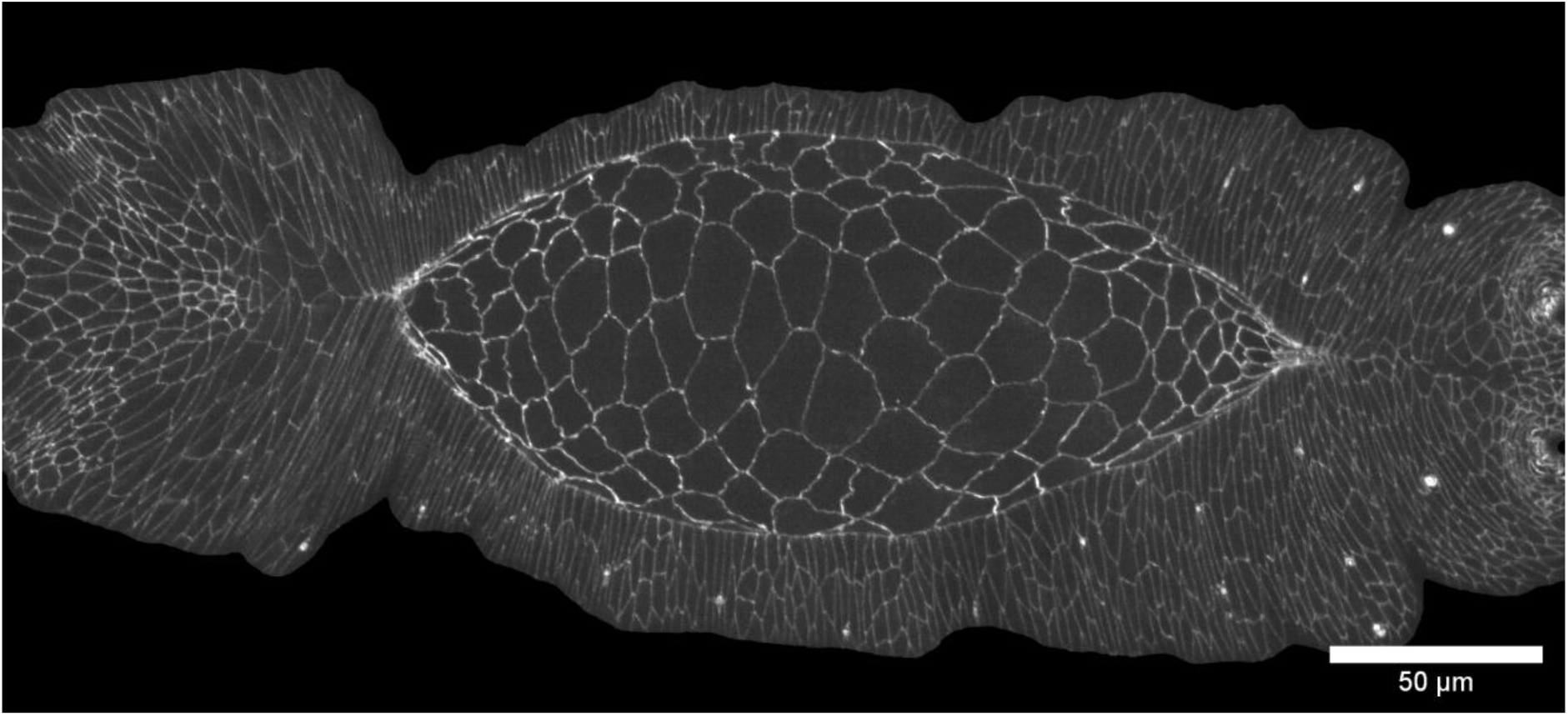
DeepProjection result of time-lapse recording of dorsal closure. Frame interval 15 s. Download: https://e.pcloud.link/publink/show?code=kZM7fJZOHPWjuH9iUpz8fS8uIp2OmHMwRyy

## Notes

### Competing Interest Statement

The authors have declared no competing interest.

